# *Pseudocapillaria tomentosa, Mycoplasma* spp., and intestinal lesions in experimentally infected zebrafish *Danio rerio*

**DOI:** 10.1101/2020.11.08.373357

**Authors:** Michael L. Kent, Elena S. Wall, Sophie Sichel, Virginia Watral, Keaton Stagaman, Thomas J. Sharpton, Karen Guillemin

## Abstract

Intestinal neoplasms and preneoplastic lesions are common in zebrafish research facilities. Previous studies have demonstrated that the neoplasms are caused by a transmissible agent, and two candidate agents have been implicated: a *Mycoplasma* sp. related to *M. penetrans* and the intestinal parasitic nematode, *Pseudocapillaria tomentosa*, and both agents are common in zebrafish facilities. To elucidate the role of these two agents in the occurrence and severity of the neoplasm and other intestinal lesions, we conducted two experimental inoculation studies. Exposed fish were examined at various time points over an 8 mo. period for intestinal histpathologic changes and the burden of *Mycoplasma* and nematodes. Fish exposed to a *Mycoplasma* isolate from zebrafish were associated with preneoplastic lesions. Fish exposed to the nematode alone or with the *Mycoplasma* isolate developed severe lesions and neoplasms. Both inflammation and neoplasm scores were associated with an increase in *Mycoplasma* burden. These results support the conclusions that *P. tomentosa* is a strong promoter of intestinal neoplasms in zebrafish, and that *Mycoplasma* alone can also cause intestinal lesions and accelerate cancer development in the context of nematode infection.

## Introduction

The zebrafish has emerged as a very important model in biomedical research^1^, second only to the mouse^2^. As with other laboratory animals, the impacts of underlying infections can seriously compromise research^**3**^. Consequently, it is important to define the etiology and pathogenesis of common diseases of laboratory zebrafish. To this end, we have previously examined data available through the Zebrafish International Resource Center (ZIRC) diagnostic program (https://zebrafish.org/health/index.php) and reviewed histopathology records of approximately 18,000 zebrafish as part of over 1,300 diagnostic cases from over 300 research laboratories spanning the periods from 2000 to 2019. These efforts reveal two common intestinal diseases: epithelial carcinomas ^**4,5**^ and infections by the nematode *Pseudocapillaria tomentosa* ^**6,7**^. Similar to other capillarid nematodes, the worm invades intestinal tissues and cause severe inflammatory changes. The intestinal neoplasms and worms have occurred in about 17% and 15% of the facilities, respectively^**8**^. The high prevalence of these intestinal neoplasms underscores the importance in determining their cause and the role the worm may play with progression of these neoplasms.

Infectious agents are being increasingly being implicated as initiators or promoters of neoplasm, and certain helminths are recognized as the cause or at least a promoter for neoplasia^9^. Kent et al.^6^ reported an association of the intestinal cancers in zebrafish with *P. tomentosa* based on review of the data from a carcinogenesis study conducted by Spitsbergen et al.^10^, where zebrafish exposed to both DMBA and *P. tomentosa* demonstrated a higher prevalence of intestinal tumors than uninfected fish exposed to this carcinogen. We recently conducted a retrospective study of the ZIRC database, and found a strong statistically significant association of the occurrence of the worms, by both case and submitting laboratories, with the cancers^8^. However, the worm is unlikely the primary cause because amongst the laboratories with the cancers, the worms were absent in 70% of the fish. In addition, Burns et al. ^11^ showed that the tumors could be readily transmitted in the absence of the worm by water borne exposure after about 9 mo., and the disease was associated with a specific *Mycoplasma* strain related to *M. penetrans*. Following that study, Gaulke et al. ^2^ performed infection studies with *P. tomentosa* and observed development of intestinal tumors after only 3 months exposure. Microbiome profiling revealed presence of *M. penetrans* related strain in individuals with tumors. In our present investigation, we conducted two exposure experiments to elucidate the roles of *P. tomentosa* and *Mycoplasma* sp. in the intestinal neoplasm in zebrafish.

## Methods

Two separate inoculation experiments were conducted to elucidate the roles of *Mycoplasma* sp. and *P. tomentosa* in intestinal neoplasia formation in zebrafish. The *Mycoplasma* sp. was originally described associated with the tumors by Burns et al. (2018)^**11**^. The *Mycoplasma* sp. used in the present study was isolated in culture from affected fish from recipient group E of that study, and henceforth is referred to as *Mycoplasma* E. The first experiment (Experiment 1) included three experimental groups; exposed to *Mycoplasma* E, exposed to *Mycoplasma* E and *P. tomentosa*, and unexposed (controls). At that time, we did not appreciate the abundance of members of the genus *Mycoplasma* in the microbiomes of our zebrafish population. Once we became aware of the extent of these bacteria in our zebrafish population that were not intentionally inoculated with *Mycoplasma*, we conducted a second experiment to validate previous findings about the impact of *P. tomentosa* alone, in the context of our fish population’s endogenous microbiota. This experiment (Experiment 2) included two groups: *P. tomentosa* exposed and controls.

### Fish and Husbandry

Both experiments used AB line zebrafish from the Sinnhuber Aquatic Resource Center (SARL). Fish from this facility are free from important zebrafish pathogens since it was established in 2007, including *P. tomentosa* ^**13,14**^. Moreover, as part of their disease screening protocols, they routinely examine sentinel and retired fish from various lines by histology conducted by one of us (M.K.), and neither the intestinal neoplasm or preneoplastic lesions has been seen in over some 3,000 fish that were examined by histology from that laboratory since 2007. Our vivarium contains flow through water, derived from charcoal filtered city water. Temperature was maintained at 27-28°C, with conductivity at 115 -125 microsiemens, and at pH approximately 7.5. Light in the vivarium is provided for 14 h/day.

### Microscopic Examinations

For both experiments, multiple time points over several months were surveyed, with about 6 fish from each replicate tank sampled per time point (Fig. 1, Table 1). Moribund fish were included in histological analysis (Table 1), whereas as those fish that died were not examined because rapid post-mortem autolysis made them unsuitable for histologic examination. Fish were euthanized by hypothermia^**15**^, the fish were gavaged with 4% paraformaldehyde^16^, and incision was made were made in the flanks near the abdomen, and then the fish were preserved in the fixative. Fish were then processed into paraffin blocks using standard methods, and about 10 serial slides were prepared and three ribbons were mounted on each slide, resulting in about 15 fish sections/slide. Alternate sections were stained with hematoxylin and eosin, and evaluated by one of us (M.K).

**Table 1.** Two trials of zerbafish exposed to Mycoplasma sp. and *Pseudocapillaria tomentosa*. Trial 1 had 3 groups, *Mycoplasma sp. exposed (*Myco*), Mycoplasma* with *Pseudocapillaria tomentosa* (Myco+W); Trial 2, 2 groups, exposed to *P. tomentosa* (Worm) and Controls. WP = weeks post exposure, W = Worms (*P. tomentosa*), M = *Mycoplasma*. NA = Not Applicable. Note for controls these data are for corresponding groups within the trial. Severity worm infection, dysplasia/hyperplasia or inflammation (Inflam) are scored 0-3. Neoplasia is positive (+) or negative (0). Prev = prevalence. Dysplasia severity (Dys-Serv) and Inflammation Severity (Inflam-Sev) is the additive score from all fish in the sample. * = moribund fish.

| Experiment 1 |  |  |  |  |  |  |  |  |  |  |
| --- | --- | --- | --- | --- | --- | --- | --- | --- | --- | --- |
| Treatment | WP-M | WP-W | Tank # | Sample size | W-Prev | Dys-Prev | Dys-Sev | Inflam-Prev | Inflam-Sev | Tumors |
| Control | 9 | 3 | 1 | 6 | 0 | 0 | 0 | 0 | 0 | 0 |
|  |  |  | 2 | 6 | 0 | 0 | 0 | 0 | 0 | 0 |
|  |  |  | 3 | 6 | 0 | 0 | 0 | 0 | 0 | 0 |
|  |  |  | 4 | 6 | 0 | 0 | 0 | 0 | 0 | 0 |
| Control | 15 | 9 | 1 | 6 | 0 | 0 | 0 | 0 | 0 | 0 |
|  |  |  | 2 | 6 | 0 | 0 | 0 | 0 | 0 | 0 |
|  |  |  | 3 | 6 | 0 | 0 | 0 | 0 | 0 | 0 |
|  |  |  | 4 | 6 | 0 | 0 | 0 | 0 | 0 | 0 |
| Control | 26 | 19 | 1 | 6 | 0 | 0 | 0 | 0 | 0 | 0 |
|  |  |  | 2 | 6 | 0 | 0 | 0 | 0 | 0 | 0 |
|  |  |  | 3 | 6 | 0 | 0 | 0 | 0 | 0 | 0 |
|  |  |  | 4 | 6 | 0 | 0 | 0 | 0 | 0 | 0 |
| Control | 32 | 25 | 1 | 6 | 0 | 0 | 0 | 0 | 0 | 0 |
|  |  |  | 2 | 6 | 0 | 0 | 0 | 0 | 0 | 0 |
|  |  |  | 3 | 6 | 0 | 0 | 0 | 0 | 0 | 0 |
|  |  |  | 4 | 6 | 0 | 0 | 0 | 0 | 0 | 0 |
| Myco | 9 | 3 | 1 | 6 | 0 | 2 | 3 | 1 | 1 | 0 |
|  |  |  | 2 | 6 | 0 | 1 | 1 | 2 | 2 | 0 |
|  |  |  | 3 | 6 | 0 | 0 | 0 | 0 | 0 | 0 |
| Myco | 15 | 9 | 1 | 6 | 0 | 2 | 2 | 0 | 0 | 0 |
|  |  |  | 2 | 6 | 0 | 2 | 2 | 1 | 1 | 0 |
|  |  |  | 3 | 6 | 0 | 1 | 1 | 0 | 0 | 0 |
| Myco | 26 | 20 | 1 | 6 | 0 | 2 | 2 | 0 | 0 | 0 |
| Myco | 32 | 25 | 2 | 6 | 0 | 2 | 2 | 0 | 0 | 0 |
|  |  |  | 3 | 6 | 0 | 2 | 2 | 0 | 0 | 0 |
| Myco | 32 | 24 | 1 | 5 | 0 | 1 | 2 | 1 | 1 | 0 |
|  |  |  | 2 | 5 | 0 | 1 | 2 | 0 | 0 | 0 |
|  |  |  | 3 | 5 | 0 | 2 | 2 | 2 | 2 | 0 |
| Myco | 35 | 27 | 1 | 5 | 0 | 0 | 0 | 0 | 0 | 0 |
|  |  |  | 2 | 5 | 0 | 3 | 3 | 0 | 0 | 0 |
|  |  |  | 3 | 4 | 0 | 2 | 2 | 0 | 0 | 0 |

| Experiment 1 (continued) |  |  |  |  |  |  |  |  |  |  |
| --- | --- | --- | --- | --- | --- | --- | --- | --- | --- | --- |
| Treatment | WP-M | WP-W | Tank # | Sample size | W-Prev | Dys-Prev | Dys-Sev | Inflam-Prev | Inflam-Sev | Tumors |
| Myco+W | 9 | 3 | 1 | 6 | 4 | 4 | 8 | 5 | 7 | 0 |
|  |  |  | 2 | 6 | 5 | 5 | 7 | 5 | 9 | 0 |
|  |  |  | 3 | 5 | 3 | 3 | 3 | 3 | 3 | 0 |
| Myco+W | 15 | 9 | 1 | 6 | 6 | 3 | 9 | 5 | 11 | 0 |
|  |  |  | 2 | 6 | 5 | 4 | 7 | 3 | 7 | 0 |
|  |  |  | 3 | 6 | 5 | 5 | 8 | 4 | 7 | 0 |
| Myco+W | 17 | 11 | 1* | 1 | 1 | 1 | 3 | 1 | 2 | 1 |
| Myco+W | 18 | 12 | 3* | 1 | 1 | 1 | 1 | 1 | 3 | 0 |
| Myco+W | 19 | 13 | 2* | 1 | 1 | 1 | 3 | 1 | 3 | 1 |
| Myco+W | 22 | 16 | 1* | 1 | 1 | 1 | 2 | 1 | 2 | 1 |
| Myco+W | 23 | 18 | 2* | 1 | 1 | 1 | 2 | 1 | 2 | 0 |
| Myco+W | 24 | 18 | 3* | 1 | 0 | 1 | 3 | 1 | 3 | 1 |
| Myco+W | 26 | 20 | 1 | 5 | 5 | 5 | 15 | 5 | 15 | 4 |
|  |  |  | 2 | 5 | 5 | 5 | 10 | 5 | 11 | 2 |
|  |  |  | 3 | 5 | 4 | 4 | 10 | 4 | 10 | 2 |
| Myco+W | 47 | 41 | 1* | 1 | 1 | 1 | 3 | 1 | 3 | 0 |
| Myco+W | 53 | 47 | 1 | 4 | 3 | 3 | 6 | 4 | 5 | 0 |
|  |  |  | 2 | 2 | 1 | 0 | 0 | 1 | 2 | 0 |
|  |  |  | 3 | 2 | 2 | 2 | 5 | 2 | 4 | 1 |

| Experiment 2 |  |  |  |  |  |  |  |  |  |  |
| --- | --- | --- | --- | --- | --- | --- | --- | --- | --- | --- |
| Treatment | WP-M | WP-W | Tank # | Sample size | W-Prev | Dys-Prev | Dys-Sev | Inflam-Prev | Inflam-Sev | Tumors |
| Control | NA | 3 | 1 | 6 | 0 | 0 | 0 | 0 | 0 | 0 |
|  |  |  | 2 | 6 | 0 | 0 | 0 | 0 | 0 | 0 |
| Control | NA | 8 | 1 | 6 | 0 | 0 | 0 | 0 | 0 | 0 |
|  |  |  | 2 | 6 | 0 | 0 | 0 | 0 | 0 | 0 |
| Control | NA | 29 | 1 | 6 | 0 | 0 | 0 | 0 | 0 | 0 |
|  |  |  | 2 | 6 | 0 | 0 | 0 | 0 | 0 | 0 |
| Worms | NA | 3 | 1 | 6 | 6 | 5 | 8 | 4 | 5 | 0 |
|  |  |  | 2 | 6 | 3 | 3 | 3 | 3 | 3 | 0 |
| Worms | NA | 9 | 1 | 6 | 2 | 2 | 2 | 2 | 3 | 0 |
|  |  |  | 2 | 5 | 0 | 0 | 0 | 1 | 1 | 0 |
| Worms | NA | 16 | 1 | 7* | 6 | 7 | 16 | 6 | 14 | 1 |
|  |  |  | 2 | 4 | 4 | 4 | 8 | 4 | 8 | 0 |
| Worms | NA | 30 | 1 | 3 | 2 | 3 | 5 | 3 | 3 | 0 |
|  |  |  | 2 | 2 | 2 | 2 | 2 | 2 | 4 | 1 |

**Figure 1.**
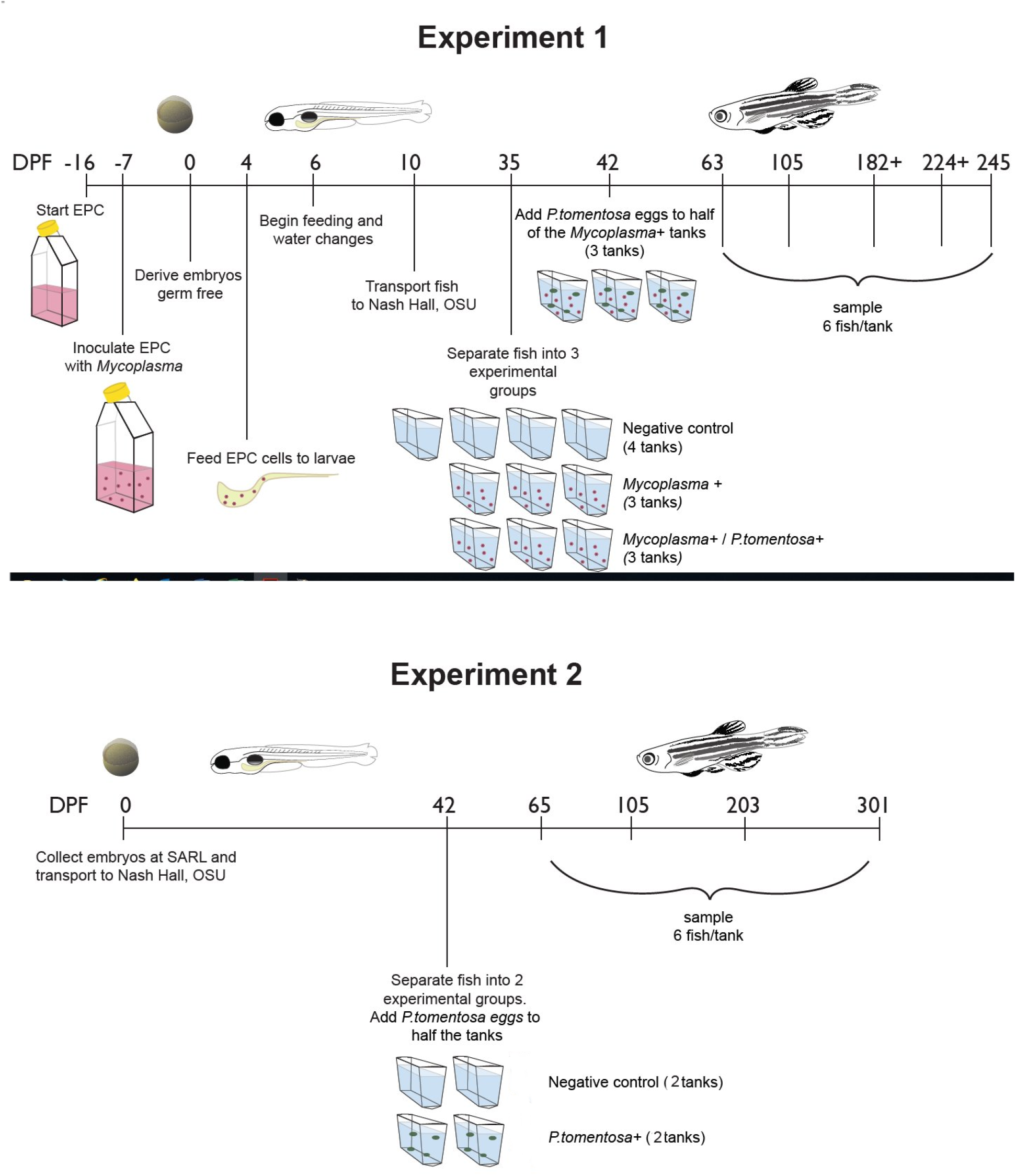
Time line for exposure and sampling of zebrafish; Experiment 1, exposed to either Mycoplasma E, Mycoplamsa E with P. *tomentosa* or Negative Controls and Experiment 2, *P. tomentosa* and Negative Controls. DPF = Days post fertilization, EPC cells = EPC cells previously infected with *Mycoplasma* E.

Each fish was scored for hyperplasia, dysplasia, neoplasia of the epithelium, and inflammation of the lamina propria as described previously^**12**^. Scores ranged from zero (absent) to three (severe) (See Fig 2). For proliferative lesions, those with neoplasia were scored as 3 as they all also had preneoplastic changes (hyperplasia or dysplasia). Severity scores at each sample time for each tank were calculated by adding all scores together and dividing the results by number of fish in the sample (Table 1). Our scoring system for these changes, 0-3, is not “ordinal”, in that fish with a score of 3 have severe lesions that were consistently far greater than 3 times as severe as a score of 1.

**Figure 2.**
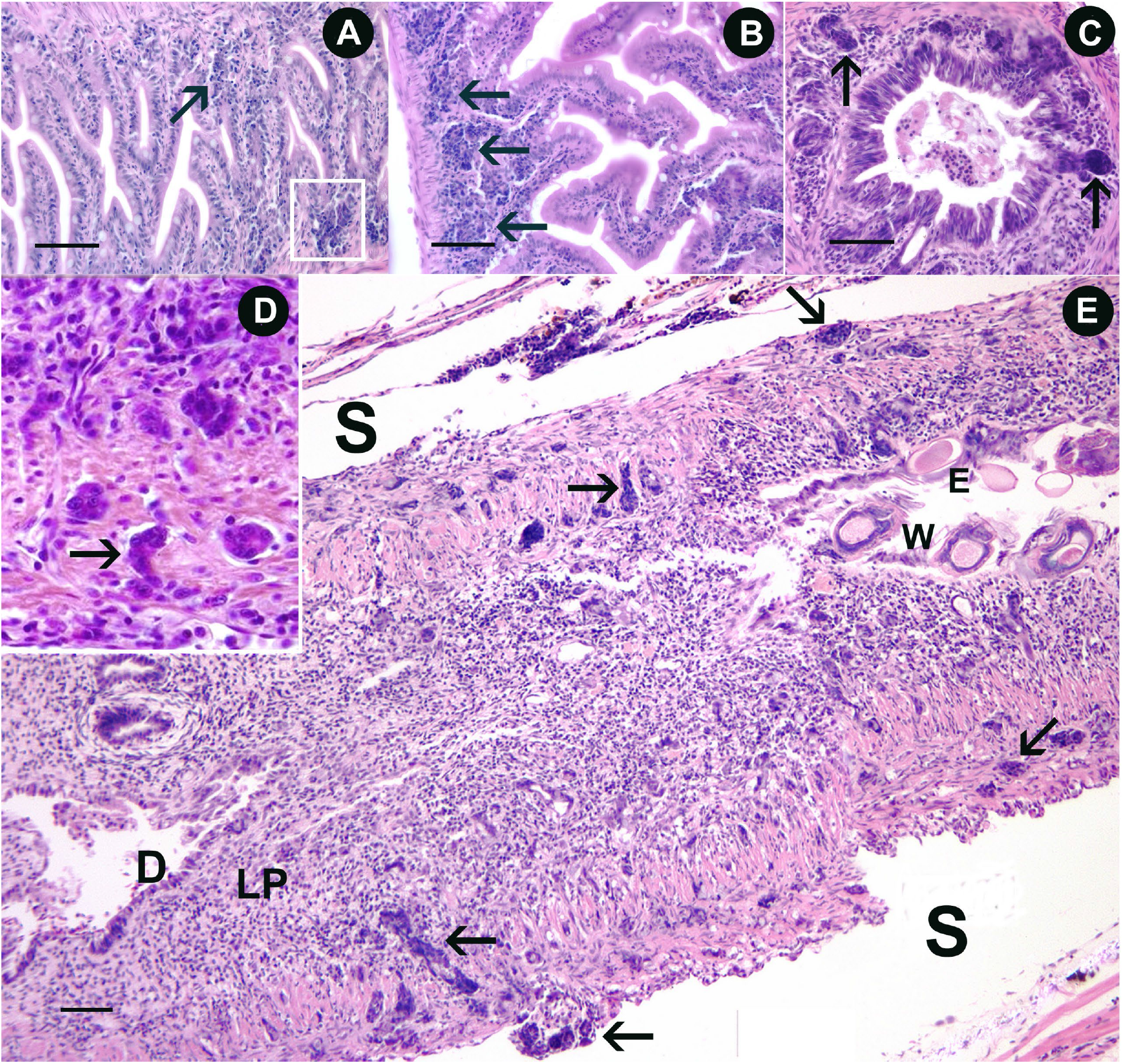
Intestine histology. Hematoxylin and eosin. Bar = 50 µm. A, B. Fish exposed to *Mycoplasma* E (Exp. 1); arrows and square = region of hyperplasia/dysplasia. Note loss of bipolar, columnar appearance of epithelial cells. C, D. Intestinal neoplasia in a fish exposed to *Mycoplasma* E and *P. tomentosa*. Arrows = pegs or nests of neoplastic epithelial cells in lamina propria and (in D) nests within the muscularis. E. Low magnification of fish exposed to *Mycoplasma* E and *P. tomentosa* (W). The epithelium (D) is severely dysplastic, flatten or absent in some regions. Laminia propria (LP) exhibits severe, chronic inflammation and nest of neoplastic occur throughout the muscularis (arrows) and into the serosa/coelom (S).

#### Fluorescent in situ Hybridization (FISH)

Unstained sections from selected fish were processed for FISH following the protocol described in our previous studies ^6^. Slides were selected from the various groups, control fish as well as exposed fish, with or without lesions by M.K. These slides were then processed for FISH and examined by one of us (E.W.) with no knowledge of the fish’s status by groups or lesions.

Slides with paraffin embedded sections were deparaffinized with Clear-Rite 3 (Richard-Allan Scientific, Kalamazoo, MI) and rehydrated through decreasing concentrations of ethanol into sterile ddH_2_0 following our general laboratory protocol ^**16**^. Prewarmed (70°C) hybridization buffer (20 mM Tris-HCl (pH 7.4), 0.9 M NaCl, 0.1% SDS, 35% v/v formamide) was applied to slides to equilibrate sections for 1 minute and then removed. Prewarmed (70°C) hybridization buffer mixed with probes at a working concentration of 25 pmol were then applied to equilibrated slides and a glass coverslip was placed on top to disperse probe mix and cover all sections. Slides were placed in a light excluding staining tray (with sterile water to maintain humidity) and incubated overnight at 48°C. After incubation, slides were soaked for 10 minutes at 48°C in sterile prewarmed 1xPBS, cover slips were removed, and slides were subsequently soaked two more times in fresh sterile 1xPBS for 10 minutes each, gradually bringing slides and PBS to room temperature. Slides were mounted with VECTASHIELD Antifade Mounting Medium with DAPI (Vector Laboratories, Burlingame, CA) and sealed with a coverslip. Slides were assessed and imaged using a Nikon Eclipse Ti inverted microscope equipped with an Andor iXon3 888 camera.

Sections were processed as described above with a mixture of *Mycoplasma* genus specific oligonucleotide probes to 23s rDNA with Cy3 attached to 5′ and 3′ ends (Mycoplasma_295-3: AAGGAACTCTGCAAATTAACCCCGTA; Mycoplasma_295-4: AAGGAACTCTGCAAATTCATCCCGTAAG). We selected these 23s rDNA probes because they successfully detected *M. penetrans* in samples using a DNA microarray^18^. FISH using these probes showed no showed no evidence of *Mycoplasma* spp. in the negative control fish. As a positive control to visualize all bacteria, including *Mycoplasma*, a mixture of eubacterial oligonucleotide probes with fluorescein attached to 5′ and 3′ ends were used (Eub338-I: GCTGCCTCCCGTAGGAGT; Eub338-II: GCTGCCACCCGTAGGTGT: and Eub338-III GCTGCCACCCGTAGGTGT) ^**17**^ As a positive control to visualize all bacteria, including *Mycoplasma*, a mixture of eubacterial oligonucleotide probes with fluorescein attached to 5′ and 3′ ends were used (Eub338-I: GCTGCCTCCCGTAGGAGT; Eub338-II: GCTGCCACCCGTAGGTGT: and Eub338-III GCTGCCACCCGTAGGTGT) ^**18**^. DAPI was used to visualize DNA in eukaryotic and some prokaryotic cells (Integrated DNA Technologies Coralville, IA) (http://www.microbial-ecology.net/probebase).

Sections were graded blindly with no knowledge of group status or histology. However, slides were selected for FISH testing that had a large amount of intestinal tissue based on observations from adjacent slides stained with H&E. The entire slide was evaluated based on a 0-5 scale by identifying areas within the intestines that were labelled with both *Mycoplasma* and eubacterial probes (Fig. 3). Scoring was as follows: 0 = no double positive signal in any sections on a slide; 1 = 1-2 puncta (particles), or 1-2 aggregates; 2 = 3-5 particles, or 3-5 aggregates; 3 = most, or every, section on slide has 1-2 particles or clumps; 4 = most, or every, section on slide has 3-4 particles or clumps; 5 = Most, or every, section on slide has 5-10 particles or clumps and almost all or all sections has >10 particles or clumps per section. Scores for each fish were then recorded in Table 2, with a total score adding particles and aggregates.

**Figure 3.**
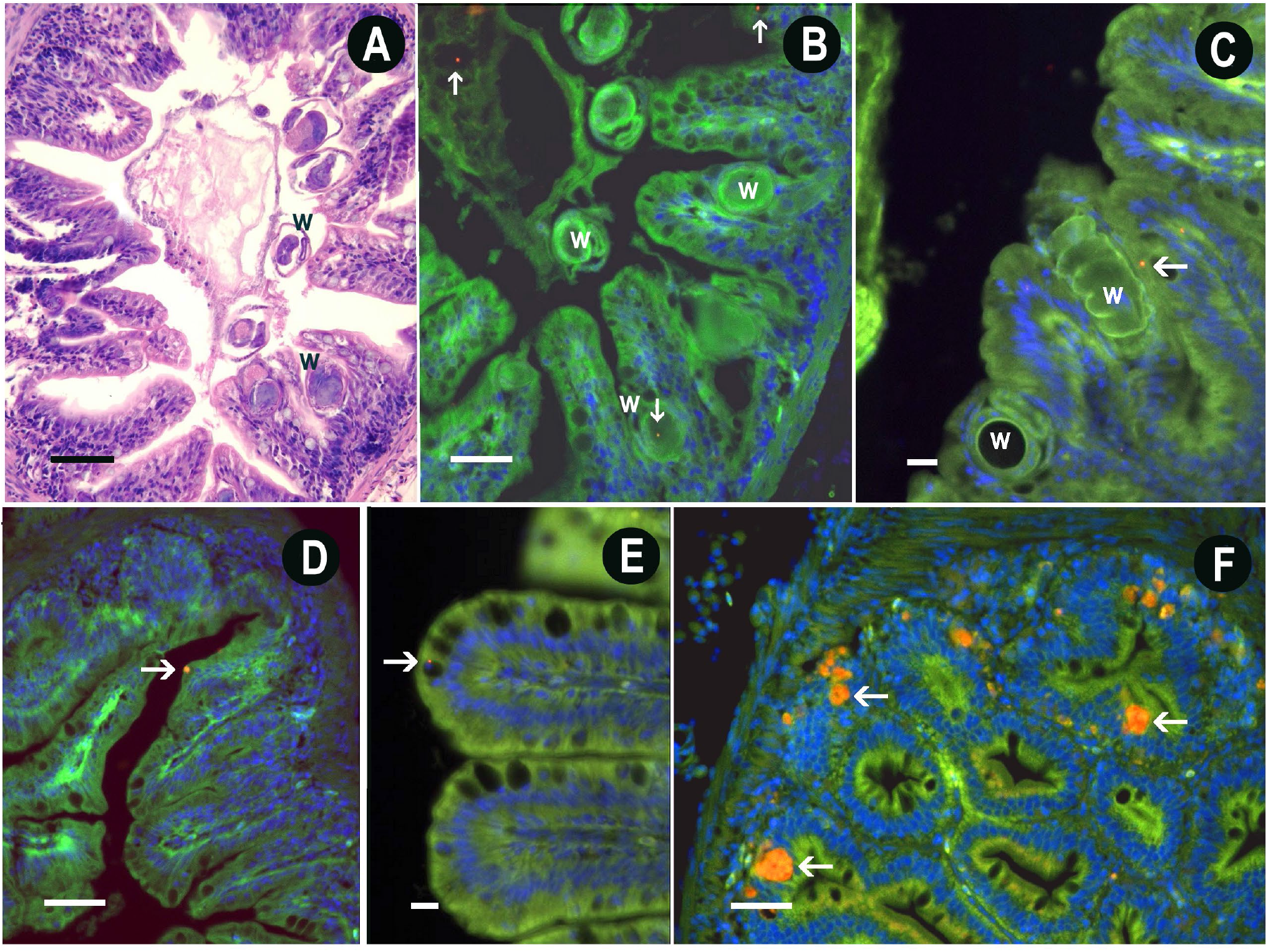
Intestines with *Mycoplasma*, and *P. tomenstosa* stained with either FISH or H&E. Bars = 50 µm for A, B, D, F = 50 µm; 10 µm for C and E. W = worms, Arrows = Red particles or aggregates, positive for *Mycoplasma*. A,B) adjacent slides stained with either H&E or FISH. Cross sections of worms either in the epithelium or in lumen. B) Particles, including one within a worm. c) FISH showing worms with adjacent particle. d) Small aggregate, intracellular at tip of epithelial cell e) Intracellular particle adjacent to goblet cell. e) Multiple intercellular aggregates.

**Table 2.**
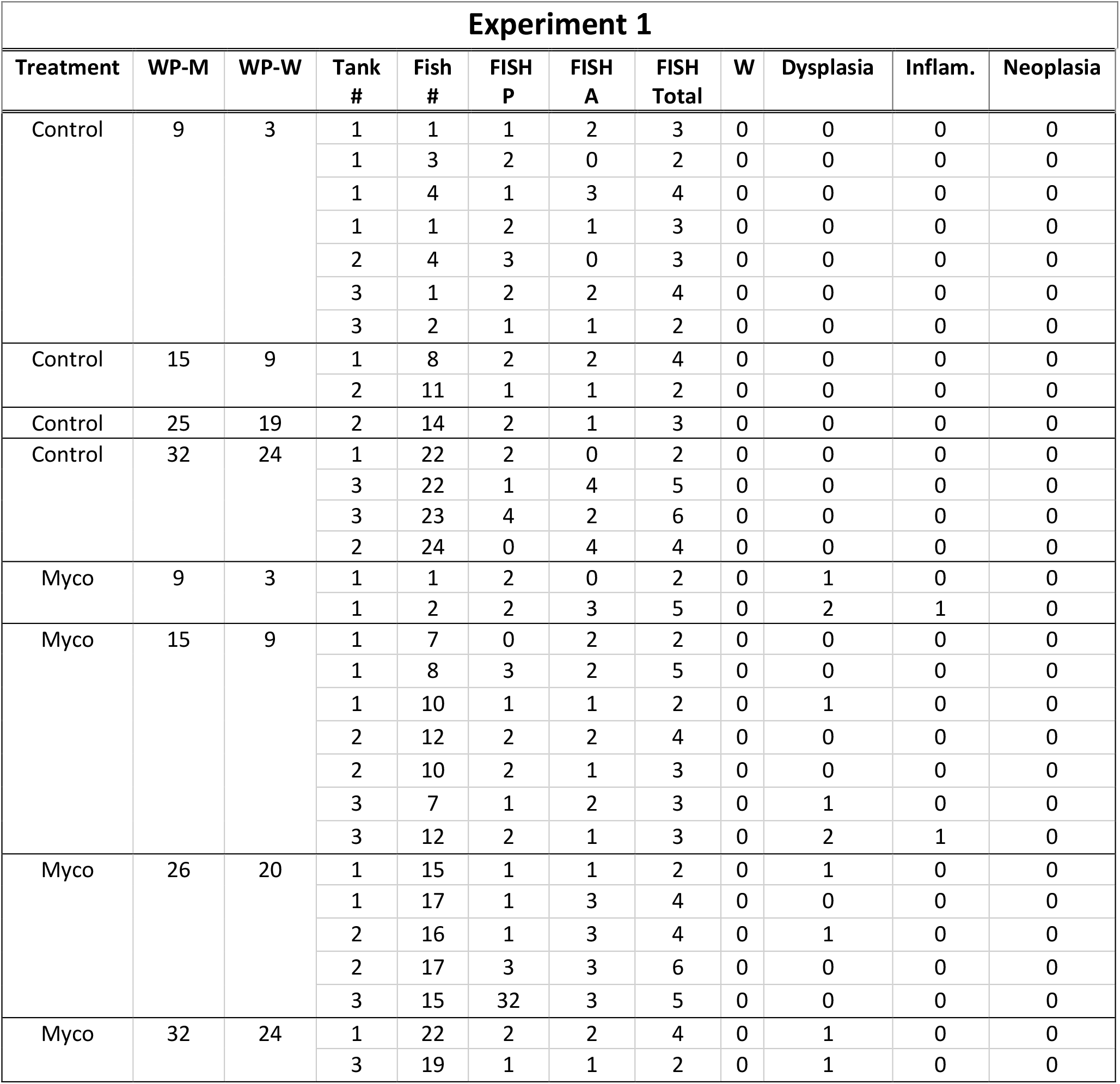
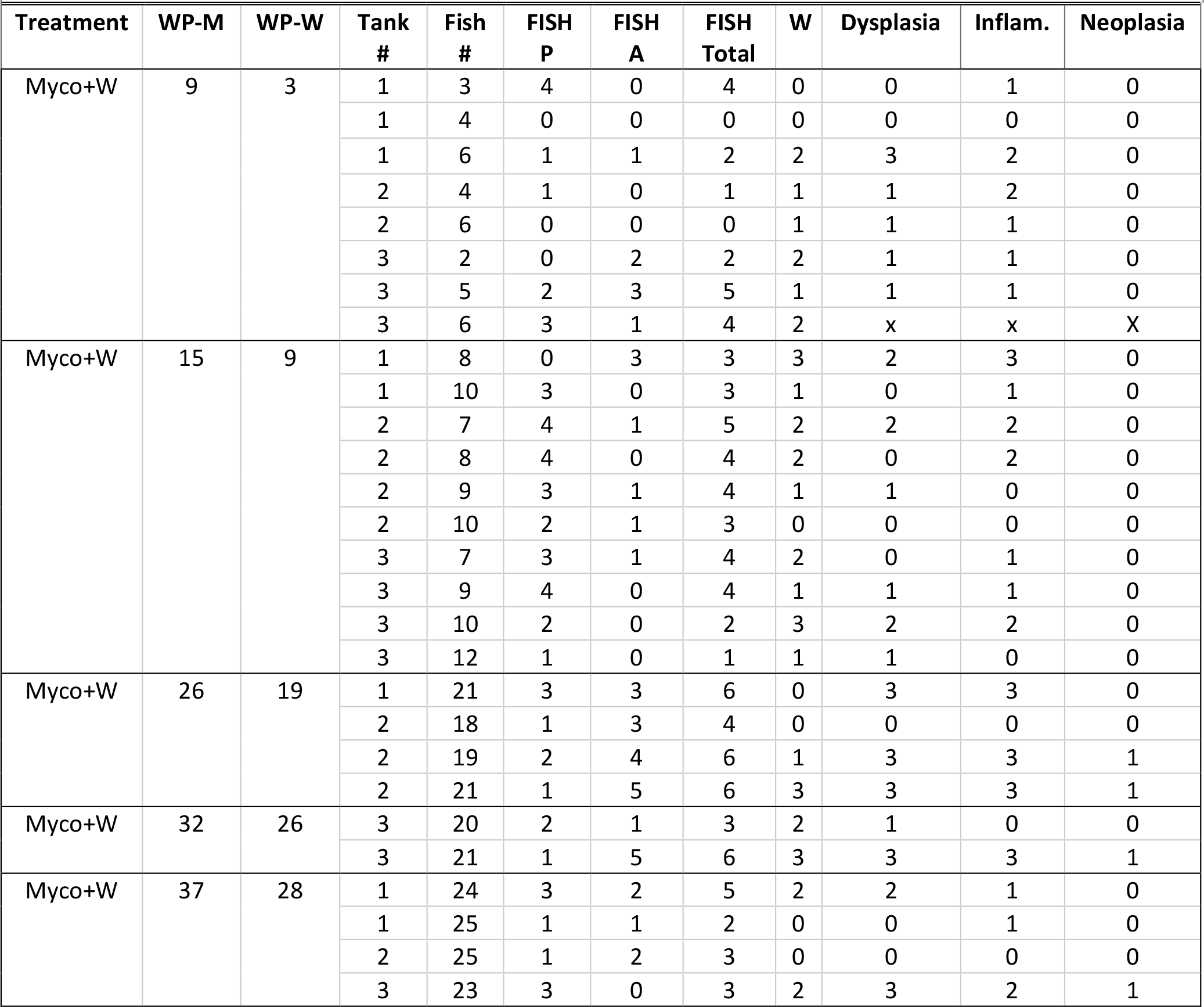
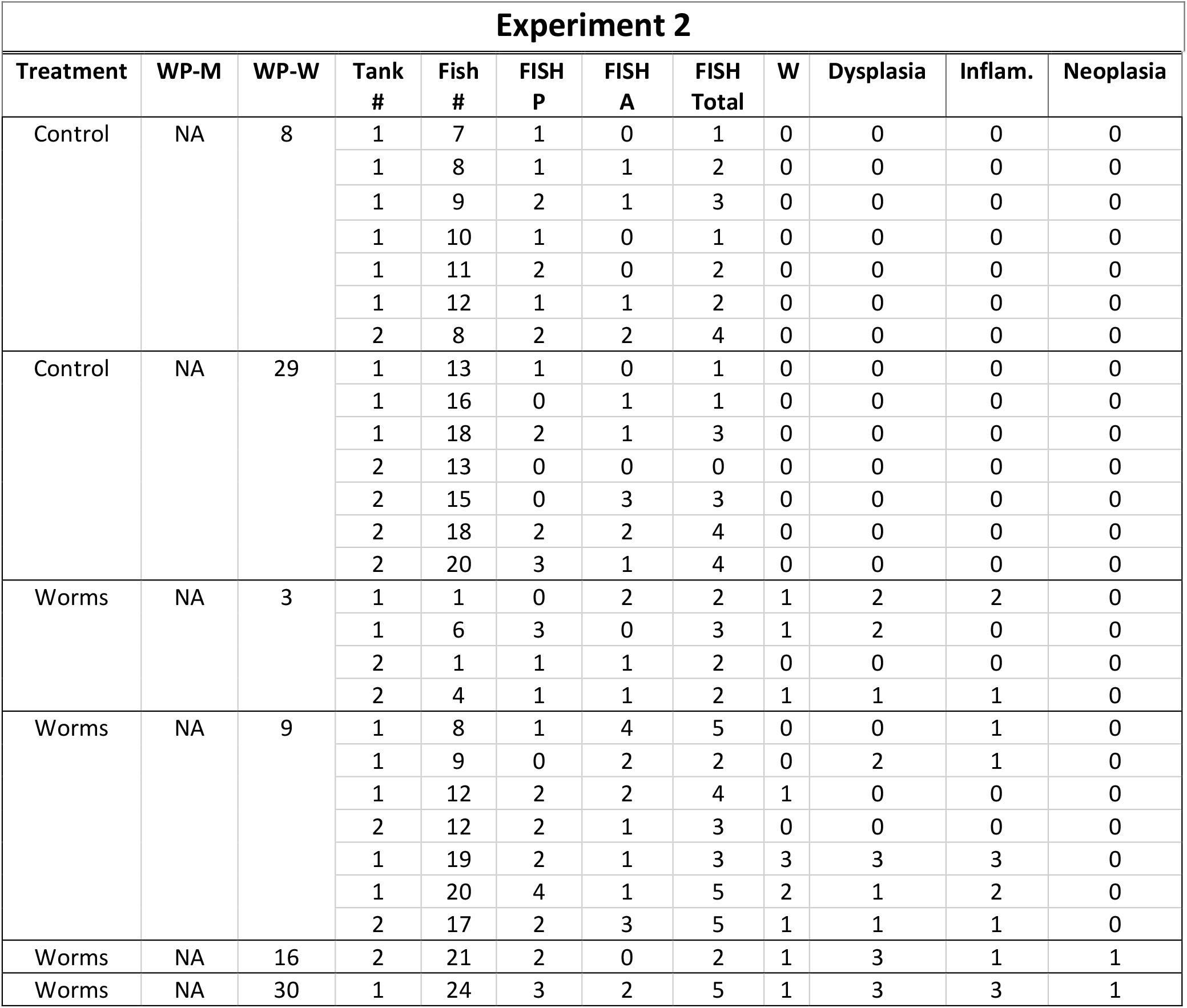
*Mycoplasma* in zebrafish intestines examined with fluorescent in situ hybridization (FISH) genus specific probe in two in vivo trials. Treatment regimens (Treat) are C = Control (no exposure), M = *Mycoplasma* exposure, W = worm exposure. WP = weeks post exposure, W = Worms (*Pseudocapillaria tomentosa*), M = Mycoplasma. WP = weeks post exposure. Note for controls these data are for corresponding groups within the trial. Fish # = individual fish number. FISH results are presented as individual particles or as aggregates of presumptive *Mycoplasma* cells. Total = Addition of scores for both particles (P) and aggregates (A). Severity worm infection, dysplasia/hyperplasia or inflammation are scored 0-3. Neoplasia is positive (+) or negative (0).

### Statistics

Statistical analyses were conducted in *R* (R Core Team 2020). To test the association between FISH and histological scores, the following procedure was conducted. We built ordinal logistic regression models of the ordered categorical histological score data using the function *polr* from the *MASS* package ^19^. Though we note that ordinal regression may underestimate histological variation given that differences in histological score categories increase along the scoring range. We started with a base model that only included Experiment ID and built increasingly complex models by adding parameters to this base model, which enabled us to determine whether the additional parameters better explained the variation in either FISH or histological scores than Experiment ID did alone. Our progressive model construction approach added parameters one by one and included interaction terms in the most complex models. We sequentially applied likelihood ratio tests to these models using the *anova* function from the base *stats* package to select the most optimal model. For all histology measures, optimal models were consistently identified as the following when considering clump scores:

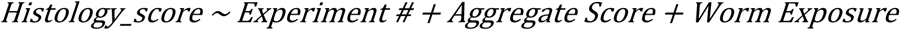

Moreover, all histology measures manifested a consistent optimal model when considering particle scores:

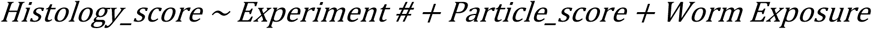

However, while these were determined to be optimal models, not all parameters in these models were necessarily significantly associated with the histological response variable. Using the selected model, we calculated odds ratios for each term and estimated their 95% confidence intervals (function *confint* in the base *stats* package). Terms with confidence intervals that overlapped zero were considered not significant and therefore not included in the reported table (Table 3). All other statistical comparisons were conducted using the two-sample Wilcoxon rank sum tests using the function *wilcox*.*test* from the base *stats* package.

**Table 3.**
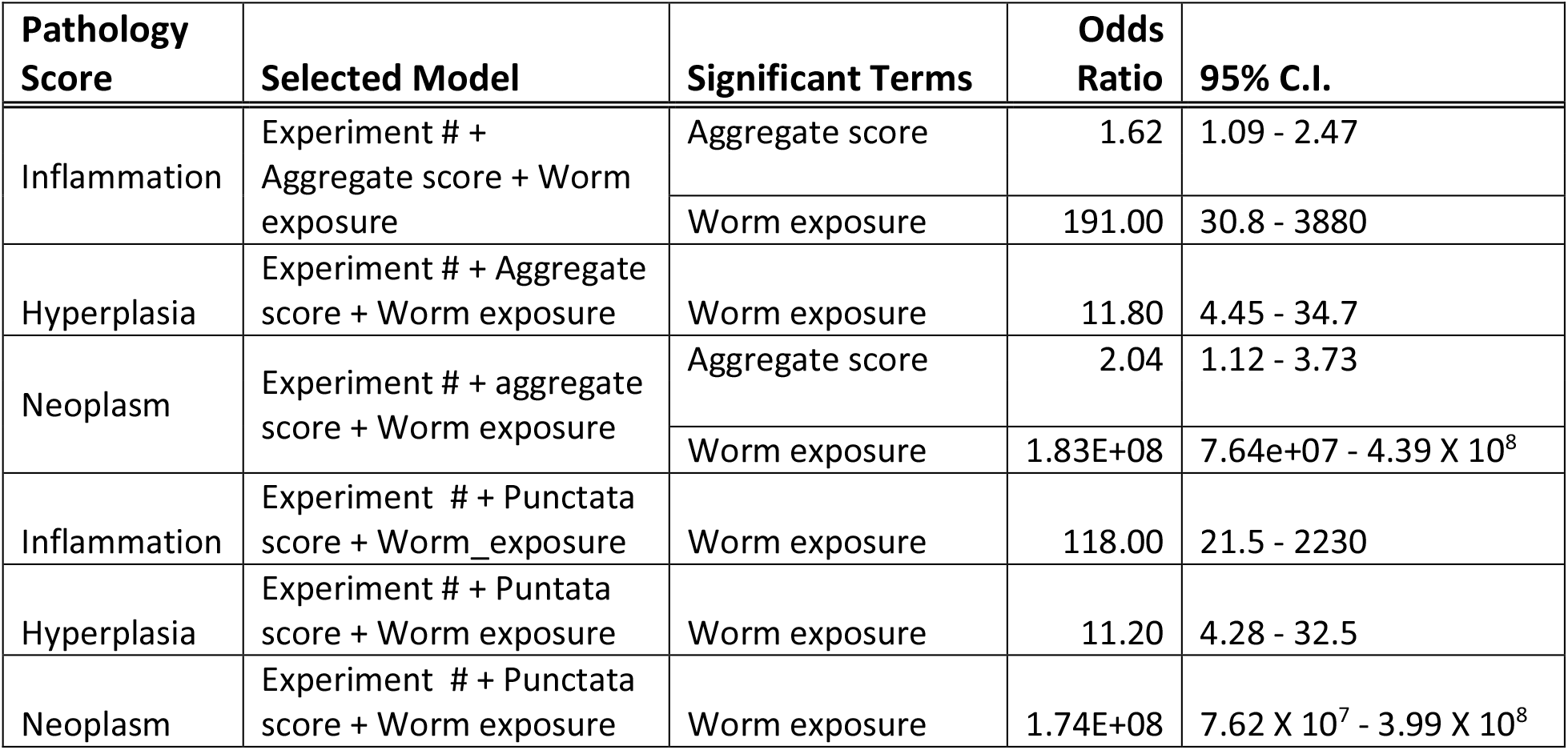
Pathologic changes treatments, FISH results (puncta or aggregates), and pathology endpoints in two zebrafish laboratory transmission experiments in which fish in which groups of fish received various treatments; *Pseudocapillaria tomentosa, Mycoplasma* E, or unexposed controls. Experiment # = Experiment 1 or 2. Hyperplasia = hyperplasia and/or dysplasia.

| Pathology Score | Selected Model | Significant Terms | Odds Ratio | 95% C.I. |
| --- | --- | --- | --- | --- |
| Inflammation | Experiment # + Aggregate score + Worm exposure | Aggregate score | 1.62 | 1.09 - 2.47 |
|  |  | Worm exposure | 191.00 | 30.8 - 3880 |
| Hyperplasia | Experiment # + Aggregate score + Worm exposure | Worm exposure | 11.80 | 4.45 - 34.7 |
| Neoplasm | Experiment # + aggregate score + Worm exposure | Aggregate score | 2.04 | 1.12 - 3.73 |
|  |  | Worm exposure | 1.83E+08 | 7.64e+07 - 4.39 X 10<sup>8</sup> |
| Inflammation | Experiment # + Punctata score + Worm_exposure | Worm exposure | 118.00 | 21.5 - 2230 |
| Hyperplasia | Experiment # + Puntata score + Worm exposure | Worm exposure | 11.20 | 4.28 - 32.5 |
| Neoplasm | Experiment # + Punctata score + Worm exposure | Worm exposure | 1.74E+08 | 7.62 X 10<sup>7</sup> - 3.99 X 10<sup>8</sup> |

### Experiment 1

The *Mycoplasma* sp. used in the present study was isolated from affected fish from our previous transmission study ^11^(Burns et al. 2018). The isolate used here was from recipient group E from that study, and hence is referred to as *Mycoplasma* E. It was isolated from zebrafish on SP4 solid media (50g/L BHI, 10g/L Peptone, 5g/L NaCl, 15g/L Yeast extract, 15g/L Agar, 200ml/L sterile heat inactivated horse serum, 20mg/L Amphotericin B, 250mg/L Ampicillin, 250mg/L Polymixin B) ^**20**^ were cut from agar and inoculated in SP4 broth (as above, without agar) for ∼7 days at 37 °C with shaking. 2 mL of culture were transferred to a cryotube and centrifuged at 16,000 x G at 4°C for 30 minutes to pellet *Mycoplasma* cells, supernatant was aspirated, and the pellet was suspended in 750uL fresh SP4 broth (room temperature) and 750uL sterile 20% glycerol. The suspension was mixed and allowed to equilibrate at room temperature for 15 minutes, it was then flash frozen in liquid nitrogen and stored at -80°C. The similarity of *Mycoplasma* E 16s rDNA to the original sequence described by Burns et al.^11^ from donor fish and *M. penetrans* from humans (ATCC 55252, GenBank JN935872.1) and the sequence from *Mycoplasma* BHJA from salmon^**21**^ GenBank # AY065998.1. 16S rRNA candidate sequences were aligned using the align.seqs command in mothur v.1.44.3 ^**22**^, and the SILVA SEED v132 database^**23**^ as a template. A distance matrix was then generated using the resulting alignment and the dist.seqs (options: calc=nogaps, output=square) command in mothur. Uncorrected pairwise distance values were subtracted from 1.0 and converted to percentages to find the percent similarity.

*Mycoplasma* spp. grow very well in cell culture^24^ and *Mycoplasma* E was also grown in cell culture, using a common fish cell line used in virology; EPC cells from fathead minnow (*Pimephales promelas*). The cells were cultured with regular media changes, at 25°C in Leibovitz L-15 media supplemented with 10% sterile Heat Inactivated Fetal Bovine Serum, Pen/Strep (1:1000 dilution, Sigma), and Gentamycin (10 µg/mL). Cells were split twice in a 7 day period, and with the second split, half the cells were suspended in the above mentioned media, while the other half were suspended in Leibovitz L-15 media supplemented with 10% sterile Heat Inactivated Fetal Bovine Serum, Amphotericin B (2.5mg/ml), Ampicillin (100mg/ml), and Polymixin B (50mg/ml), or *Mycoplasma* media. Flasks containing *Mycoplasma* media were inoculated with *Mycoplasma* E glycerol stock using wooden sticks, flasks containing regular EPC growth media were exposed to sterile wooden sticks. Inoculated and control flasks were maintained at 30°C, with regular media changes. 24 hours before inoculation into fish flasks, media was changed but all cells were washed with sterile 1XPBS and given *Mycoplasma* media. On the day of inoculation into fish cell line flasks, cells were harvested by centrifuging cells and media at 1000 x g for 5 minutes, after the cells were pelleted the media was removed and cells were suspended in sterile 1xPBS. Harvested EPC cells were counted and inoculated into each flask containing 4 days post fertilization (dpf) larvae at a density of ∼2.21 x 10^6^ EPC cells/ flask.

Figure 1 provides a time line of the experimental design for exposure to *Mycoplasma* E and *P. tomentosa*. Approximately 800 embryos were transferred to University of Oregon and bacterial loads were depleted using a previously described chorion disinfection method^25^. Surface sterilized embryos were placed in sterile T75 filtered tissue culture flasks (TPP, Trasadingen, Switzerland) containing 50mL sterile embryo medium EM at a density of ∼1 embryo/2mL. Flasks of zebrafish larvae were maintained at 28-30°C within a room with a 14hr/10hr light/dark cycle. At 4 dpf, when zebrafish first start feeding, one half of the fish were exposed to *Mycoplasma* by feeding EPC cells from infected flasks, whereas the negative controls were fed uninfected EPC cells. Starting at 6 dpf, larvae from both groups were transferred to petri dishes at density of ∼1 larvae/2mL. Larvae were fed rotifers (*Brachionus plicatilis*) and given health checks twice a day, with water changes once a day. At 10dpf, each petri dish of larvae was transferred to a fresh T75 filtered tissue culture flasks with 150mL sterile EM. Flasks we transported to Oregon State University for further maintenance. Unexposed larvae were examined by FISH using the *Mycoplasma*-specific probe.

Fish were transferred to the Kent laboratory at OSU at 10 dpf and they continued to be fed rotifers provided by the Animal Care Services, University of Oregon and Gemma Micro diet (Skretting, Wesbrook, ME) was included for 21 dpf. Fish were then feed on for the remainder of the study. Fish were held in two 9 L aquaria (exposed and controls) with a slow drip water exchange until 24 dpf. At 39 dpf (35 days post exposure to *Mycoplasma*), fish were transferred to 9L tanks and set up into the three groups as follows: 1) *Mycoplasma* E only (3 replicates tanks), 2*) Mycoplasma* with *P. tomentosa* (3 replicate tanks) and Negative Controls (4 replicate tanks). Control fish were held in a separate, adjacent room in the vivarium, but received water from the same system. Note that a fourth group, exposed to worms but not *Mycoplasma* was not included as early in the experiment we discovered that *Mycoplasma* spp. were present in controls by testing feces with the PCR test as described in Burns et al.^11^.

For worm exposures, the *Mycoplasma* E + *P. tomentosa* group was exposed to 75 larvated worm eggs/fish as described in ^26^ when the fish were 42 dpf. The nematode eggs were disinfected with 30 ppm sodium hypochlorite for 10 min, and then treated with sodium thiosulfate before exposure as previously described ^26^ to reduce concurrent bacteria from the donor fish for these parasites. Six fish from each tank were collected and examined by histology at 4 time points from 2 through 8 mo. All groups of fish in tanks examined daily, and moribund fish were euthanized and processed for histology, whereas dead fish were not analyzed due to severe post-mortem autolysis.

### Experiment 2

We conducted a second experiment following the discovery that members of the genus *Mycoplasma* were more widespread in zebrafish and occurred in Experiment 1 controls. Also, an experiment in our laboratory showed that fish exposed to *P. tomentosa* may develop tumors as soon as 3 months post exposure12. This experiment had two groups; exposed to *P. tomentosa* and negative controls, each with a replicate tank. The same population of zebrafish fish used in the present Experiment 1 were reared from embryos in the Kent laboratory at OSU. Fish were divided into 4 tanks and 2 tanks received chlorine nematode eggs at 50 eggs/fish as described above. Similar to Experiment 1, fish from each tank were examined over a 8 month period (Fig. 1, Table 1).

## Results

Fish exposed to the nematode in both experiments exhibited profound pathologic changes in the intestine, including the occurrence of neoplasms in several fish. Some fish exposed to *Mycoplasma* E alone exhibited mild to moderate hyperplastic or dysplastic changes (Fig. 2, Table 1), whereas all controls in both experiments showed no histologic changes in the intestine.

### Experiment 1

A total of 23 of 83 fish in the *Mycoplasma* E only group had mild hyperplasia or dysplasia of the intestinal epithelium, which was first observed in the 9 week post exposure (wk PE) sample. An additional 4 fish had equivocal proliferative changes in the epithelium, but they were scored as negative due to the uncertainty in diagnosis. Amongst the fish positive for epithelial changes in this group, six fish also exhibited mild, diffuse chronic inflammation of the intestinal lamina propria. No lesions were observed in the control fish. A Wilcoxon rank sum test showed that hyperplastic lesions were significantly more numerous in the fish exposed to only *Mycoplasma* E (W = 2,772; *p* = 0.0342).

With the fish exposed to both *Mycoplasma* E and the nematode, a total of 59 fish that were exposed to *Mycoplasma* E and the nematode were examined, and the nematode was observed in 91.0% of the fish (Table 1). A range of lesions from mild to severe hyperplasia and dysplasia was observed in 91.5% of the fish (Fig. 2). In addition, 6 fish had profound dysplasia of the epithelium to the extent that the intestinal plates were completely lost, and the underlying lamina propria was severely inflamed (Fig. 2E). Starting at 9 wk PE to the worm (15 wk PE to *Mycoplasma* E) 13 of 59 fish (22%) had lesions consistent with neoplasia, with “pegs” of putative neoplastic epithelial cells penetrating through the lamina propria and muscularis (Fig 2 C-E). Seven moribund fish were included in examinations, and they consistently exhibited more severe lesions and most had the neoplasm (Table 1). And additional 12 fish died in the study and were not included due to severe post-mortem autolysis. Wilcoxon rank sum tests showed that fish exposed to both *Mycoplasma* E and the nematode carried a higher prevalence of preneoplastic lesions (W = 2,556; *p* << 0.0001) as well as neoplasms (W = 4,716; *p* = 0.0029) compared to fish only exposed to *Mycoplasma* E.

Percent sequence identity of the 16S rRNA gene was used to gauge the taxonomic proximity of Mycoplasma E and the other notable *Mycoplasma* strains (Table 4). Mycoplasma E was actually more closely related to *M. penetrans* from humans (95.4%) compared to the original Burns et al.^11^ (94.0%), whereas the sequence from Burns et al.^11^, was identical to the sequence from salmon^21^

**Table 4.** Percent identity of 16s rDNA sequence amongst Mycoplasma; *Mycoplasma penetrans* from human, the original zebrafish isolate from Burns et al. ^11^ (Myco Burns), Myco E, and the *Mycoplasma* sequence from salmon from Hoblen et al.^21^(Myco Salmon). Diagonal is the number of base pairs used in analysis.

|  | Myco E | Myco Burns | *M. penetrans* | Myco Salmon |
| --- | --- | --- | --- | --- |
| Myco E | 670 bp | 94.0 | 95.4 | 90.9 |
| Myco Burns |  | 200 bp. | 94.8 | 100 |
| *M. penetrans* |  |  | 686 bp | 92.7 |
| Myco Salmon |  |  |  | 351 bp |

### Experiment 2

Fish exposed to the nematode that were examined (n=32) showed 64% prevalence of infection by the worm. As in Experiment 1, these fish exhibited a range of lesions from mild to severe hyperplasia and dysplasia (81%), as well as inflammation of the lamina propria (78%) (Table 1; Fig. 4). Two fish were diagnosed with the intestinal neoplasm at 23 and 37 wk PE. A total of 8 fish amongst both tanks died throughout the experiment, starting 8 wk PE to the worm, but these fish were not examined. As in Experiment 1, no lesions were observed in 36 control fish. Fish exposed to the nematode carried as significantly greater number of lesions as compared to these control fish as determined by a Wilcoxon rank sum test (Wilcoxon rank sum test: W = 490; *p* << 0.0001).

**Figure 4.**
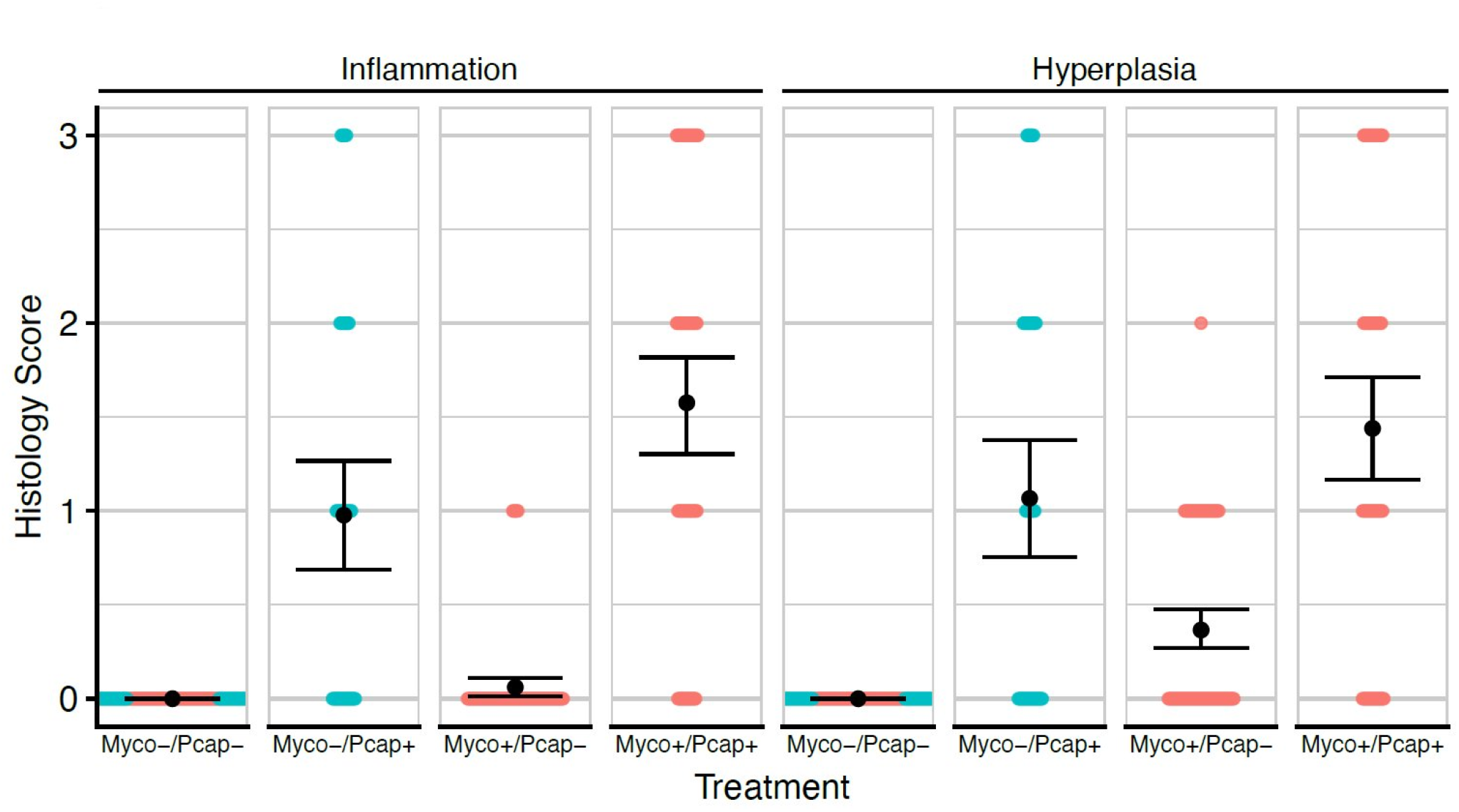
Histology scores (Inflammation or Hyperplasia/Dysplasia) for the two exposure experiments. Blue = Experiment 1, Orange = Experiment 2. X-axis indicates exposure condition: untreated (Myco-/Pcap-), exposed to *Mycoplasma* E (Myco+/Pcap-), exposed to only to *P. tomentosa* (Myco-/Pcap+), or exposed to both pathogens (Myco+/Pcap). Black dots indicate means for the scores for the given treatment and the black error bars indicate the bootstrapped 95% C.I.s for the means.

### FISH

Slides from zebrafish from all groups examined by FISH, including controls from both experiments, were positive for *Mycoplasma* spp. (Table 2). *Mycoplasma* signal often occurred associated with the intestinal epithelium as individual puncta (corresponding in size to single bacterial cells) or aggregates of positive (red) staining (corresponding in size to of an aggregate of many bacterial cells) (Fig 2, 3). In addition, *Mycoplasma* FISH signal was observed within worms (Fig. 3b), but not within worm eggs. The approximate abundance of *Mycoplasma* sp. based on FISH was similar amongst all groups, but the distribution of the *Mycoplasma* was associated with pathology. Specifically, increased abundance of aggregates was modestly but significantly associated with both inflammation and neoplasm scores (Table 3). No such associations were observed between the abundance of puncta and pathology (Table 3).

## Discussion

Infectious agents are increasingly being associated with neoplasia ^**27,28**^. Development of cancer is usually considered to require at least two mutations in the host cell of origin, and often results from multifactorial exposures and mutations. Thus clarification of the precise role of underlying chronic infections and concurrent inflammation, either as the initiator or a promoter, is challenging. Infectious agents, from oncogenic virus, bacteria, protozoa and helminths, are increasingly being associated with neoplasia^27^. With helminths, some well-recognized examples including *Schistomsoma haematobium* in urinary bladder cancer, and *Opisthorchus viverrini, O. felineui* and *Clonorchis sinensis* with liver cancer ^**29,30**^. Regarding intestinal neoplasia, *Schistosoma japonicum* has been linked to colorectal adenocarcinomas in humans through case-controlled and epidemiological studies^**31**^. Probably the best example of a nematode linked to cancer is *Spirocera lupi* in dogs, which penetrates the esophageal lining and sarcomas develop at the site of infections ^**32**^. Bacteria have also been associated with cancers, with *Helicobacter pylori* being the best studied example of a carcinogenic bacterium and driver of gastric cancer^**33**^. In the much more densely colonized distal gastrointestinal tract, several bacteria have been implicated in the development of colorectal cancer including *Fusobacterium nucleatum*, Enterotoxigenic *Bacteroides fragilis*, and *Escherichia coli* expressing the polyketide synthase genomic island^**34**^. Growing evidence finds associations between *Mycoplasma* spp. and cancer. Yang et al.^**35**^ reported an association between *Mycoplasma hyorhinis* and gastric and lung cancers through immunohistochemistry and PCR examinations of human cancers, and was supported by studies with mice and cell culture. There are also various lines of evidence of *Mycoplasma* spp. as a cause or exacerbating prostate cancer ^**36**^. *Mycoplasma penetrans* in particular has been shown to promote malignant transformations in the gastric epithelium of immunocompromised mice ^**37**^.

The mechanistic basis for microbial-associated carcinogenesis are likely to be diverse ^38^. In some cases, a specific microbial toxin is implicated in driving cancer formation, as in the case of the *H. pylori* translocated effector protein CagA in zebrafish^**39**^ or the DNA alkylating genotoxin colibactin of *E. coli* ^**40**^. In the case of nematode infections, specific metabolites from worms may lead to formation of DNA adducts, and thus implicating worms may actually be initiators, rather than just merely promoters, of cancer^**9**^. In *Mycoplasma* spp., Yang et al.^**35**^ identified a specific protein (p37) in *M. hyorhinis* that acted as a carcinogen and Zella et al (2018) ^**41**^ reported a *M. fermentans* DnaK with oncogenic properties. In other instances of microbial-associated cancers, specific microbes or microbial consortia promote generic carcinogenic processes, such as chronic inflammation, or inhibit protective processes such as immune clearance of cancer cells^**28**^. Chronic inflammation is a hallmark of chronic helminth infections which have been linked to cancer by various mechanisms including cytokine alterations^**29**^. In addition, liver fluke infections have been associated with increased of *H. pylori* suggesting that chronic inflammation can be caused by infection-induced changes in the host microbial ecology ^**29**^. Mycoplasmas are also well known to promote generic carcinogenic processes, such as upregulating inflammation and inhibiting p53 mediated cell cycle control and apoptosis^42^. There is another example of bacterial promoters in neoplasia with fish in research; *Mycobacterium marinum*, probably through chronic inflammation, increases the occurrence of liver cancers in a Japanese medaka (*Oryzias latipes*) exposed to benzo-a-pryene^**43**^.

Our earlier transmission experiments suggested that *Mycoplasma* spp. occurred at only low level, background in populations of zebrafish without tumors ^**11**^. The original isolate from donor fish was related to *M. penetrans*, and our subsequent study showed that more than one sequence variants form a clade of these related bacteria^**12**^. Likewise, *Mycoplasma* E used in the present study came from a recipient group of fish from our previous transmission study ^**11**^, and it showed minor sequence differences from the original description from donor fish. Gaulke et al.^**12**^ and our study here demonstrate that *Mycoplasma* spp. are more abundant in the zebrafish intestinal microbiome than we previously thought based on our earlier study^**11**^. This extends to other fishes, as Hoblen et al.^**21**^ showed that 96% of the microbiome of certain populations of wild Atlantic salmon (*Salmo salar*) was composed of *Mycoplasma penetrans*-like bacteria. Moreover, a recent study also showed that members of the Mycoplasmataceae were common in salmon^44^. Interestingly, like *Mycoplasma* E, the 16s rDNA sequence from the *Mycoplasma* sp. from salmon ^21^ was identical to the sequence from zebrafish in Burns et al^11^. Hence it would be reasonable to assign all of these isolates from zebrafish and salmon to *Mycoplasma penetrans sensu lato* given that the 16s rDNA differences between different species within the genus *Mycoplasma* are so dissimilar. Further taxonomic clarification on the relationships of isolates from fish with *M. penetrans* and related species will require analysis of more complete rDNA sequences, including more sequences from fish, and perhaps other genes.

Using our *Mycoplasma* FISH probe, which was specific only to the genus level, we observed similar levels of *Mycoplasma* amongst all zebrafish treatment groups, regardless of whether they were exposed to worms, *Mycoplasma* E or neither. In the present study when the worm was absent, only fish deliberately exposed to *Mycoplasma* E developed hyperplastic or dysplastic lesions in the epithelium. Preneoplastic changes (hyperplasia leading to dysplasia) in the epithelium, and inflammation in the lamina propria have been consistently observed in populations of zebrafish that have the neoplasm^**4**, **11**, **12**^. The earliest time in which tumors were observed following *Mycoplasma* exposure by co-habitation with infected fish was 8 months ^**11**^, and most of these tumors in zebrafish from research facilities are not seen until fish are about 1 year old ^**4**^. In contrast, in Experiment 1 almost all of the fish were collected before 8 months, and perhaps tumors would have developed without exposure to *P. tomentosa* if they had been incubated with the *Mycoplasma* E isolate for longer. Now that it is recognized that the *M. penetrans*-like bacteria are actually a clade with some variation, it is possible that amongst this clade some strains are more carcinogenic.

Evidence suggests that both *P. tomentosa* and *Mycoplasma* E enhance development of the intestinal tumors commonly seen in zebrafish, including both in vivo laboratory experiments ^**6**, **10**, **11**, **12**^ and in retrospective evaluation of hundreds of diagnostic cases submitted to the ZIRC diagnostic service^**8**^. In regards to the nematode, it enhances both the incidence to intestinal cancer in fish exposed to a chemical carcinogen ^**6,10**^ and also shortens the time to onset of the tumors^**12**^. It was unlikely that the tumors seen in fish exposed to the worm were caused by an undetected agent that came from the donor fish for the worms as we pretreated the worm eggs with chlorine at a relatively high concentration.

It is possible that *Mycoplasma* and *P. tomentosa* act synergistically to cause the lesions and ultimately tumors. Fish infected with both *P. tomentosa* and *Mycoplasma* E (in Experiment 1) had 22% tumors versus 4 % in the fish in Experiment 2 that were exposed to the worm without deliberate exposure to the *Mycopasma* E isolate. However comparison of results between the two experiments should be done with caution as fish were exposed to a lower dose of worms and a lower prevalence of infection occurred in Experiment 2. Severity of disease is often increased with co-infections by bacteria and helminths, including examples with *Mycoplasma* spp. Both *Mycoplasma penetrans* and *P. tomentosa* are closely associated with the epithelium, and we observed a signal for *Mycoplasma* inside of a worm. Capillarid nematodes and *Trichuris* spp. are taxonomically related, and both cause chronic, penetrating infections of the gastrointestinal epithelium. Infections by *Trichuris suis* in pigs has been associated with an increase in *Campylobacter* bacteria ^**45**^, and *T. suis* infections enhances the severity of infections by *Campylobacter jejuni* and associated lesions ^**46**^. Interestingly, amongst the pathological changes that increased with pigs with co-infections by the whipworm and the bacterium, hyperplasia of the epithelium was particularly more severe ^**45**^, and hyperplasia is a preneoplastic lesion that is consistently associated with the intestinal tumors in zebrafish ^**4**, **11**^. Future experiments are planned to disentangle the relationships between *Mycoplasma* spp., specifically *Mycoplasma* E and related isolates, the nematode, and the intestinal cancers.

## Acknowledgements

This study was supported in part by NIH ORIP 5 R24 OD010998 (MLK, TJS), NIHNIAID R21 AI135641 (TJS, MLK), NIH R01CA176579 (KG, MLK). National Institute of Environmental Health Sciences supported Environmental Health Sciences Center P30ES000210 provided SPF fish. The content is solely the responsibility of the authors and does not necessarily represent the official views of the NIH. We would like to thank Poh Kheng Loi of the Histology and Genetic Modification (HGeM) Core Facility at the University of Oregon for slide preparation. We thank Tristan Ursell, Oregon State University for providing space and equipment to inspect and image slides for FISH.

## Disclosure Statement

No competing financial interests exist.

## Notes

### Competing Interest Statement

The authors have declared no competing interest.

